# Specific and redundant roles for Gli2 and Gli3 in establishing cell fate during hair follicle development

**DOI:** 10.1101/2025.01.14.632991

**Authors:** Gokcen Gozum, Lisa Wirtz, Mareike Damen, Viktoria Reckert, Peter Schettina, Melanie Nelles, Hisham Bazzi, Catherin Niemann

## Abstract

Ectodermal appendage formation requires hedgehog (HH) signal reception, its conduction through the primary cilium and activation of Gli transcription factors (TF). How HH signalling induces cell-type-specific responses through Gli TF in stem cells (SCs) and their cilia-dependence remain open questions. Here, we use conditional mouse mutants to genetically dissect the roles of Gli and cilia HH mediators in the skin epithelium. Upon Gli2 depletion, hair follicle (HF) morphogenesis is delayed whereas sebaceous gland (SG) formation is enhanced, suggesting a dual role for Gli2 during appendage formation. By controlling proliferation of SG SCs, Gli2 impacts on the number and size of individual SG lobes. Whereas ablation of Gli3 shows no detectable phenotypes, HF cell fate is blocked in Gli2/3 double knockouts. In addition, loss of cilia phenocopies the loss of Gli2 but not the Gli2/3 mutants. Our study reveals that Gli3 exerts activator and cilia-independent functions in the absence of Gli2.

**Bullet points:**

- Epidermal Gli2 and Gli3 together mediate Hedgehog signalling and are essential for proper appendage formation
- Gli3 activator function compensates for the loss of Gli2 during HF morphogenesis
- Gli2 controls HF stem cell sub-compartments and SG formation
- Cilia mediate epidermal Gli2 but not Gli3 signalling in the skin epithelium

## INTRODUCTION

Deciphering the molecular and cellular networks driving the induction and maintenance of complex epithelial structures remain a major challenge in regenerative medicine. Morphogenesis of the mammalian skin involves an intricate crosstalk between different cell types and the Hedgehog (HH) signalling pathway emerged as a key player not only for the development of epithelial appendages, like hair follicles (HF) (Chiang et al. 1999; Sennett and Rendl 2012), but also for the process of HF neogenesis of injured skin (Lim et al. 2018; Sun et al. 2020; Frech et al. 2022; Liu et al. 2022). Importantly, abnormal activation of HH in epithelial cells induces cancer (Oro et al. 1997) and constitutes the main driver for basal cell carcinoma (BCC) formation in the skin, thus highlighting the necessity to better understand the cellular mechanisms of HH signalling particularly in the skin epithelium to allow for therapeutic strategies preventing disease formation.

The HH signal is translated into cell-specific responses by members of the Glioma-associated oncogene homolog (Gli) family of transcription factors (Zhang and Beachy 2023; Briscoe and Therond 2013). Upon HH ligand exposure, cells respond by post-translational processing of the Gli protein in predominantly transcriptional activators (Gli^A^). In the absence of HH ligands, Gli proteins can function as transcriptional repressors (Gli^R^) preventing the expression of HH target genes (Aza-Blanc et al. 1997; Methot and Basler 1999; Niewladomski et al. 2014). Mammalian HH signalling and Gli processing into Gli^A^ or Gli^R^ may require the primary cilium, an antenna-like protrusion present in most post-mitotic cells in the mammalian body (Nozawa et al. 2013). The movement of HH signalling intermediates into and out of the cilium is facilitated by intraflagellar transport (IFT) proteins. Defects in various components of cilia result in severe birth abnormalities termed ciliopathies that are also characterised by mis-regulation of HH target genes (Huangfu et al. 2003; Ho and Stearns 2021).

Mechanistic studies showed that in the absence of HH ligands, the HH receptor patched 1 (Ptch1) suppresses its co-receptor smoothened (Smo), HH ligand binding to Ptch1 inhibits Smo suppression, allowing it to move to the primary cilia and activate HH signal transduction, including Gli-mediated expression of HH target genes (Zhang and Beachy 2023). The disruption of the IFT complexes, including IFT88, leads to the loss of cilia and subsequently interferes with HH signalling (Huangfu et al. 2003; Houde et al. 2006). The role of primary cilia in HH signal transduction is complex and context-dependent. Given the importance of the HH-cilia signalling axis in numerous developmental processes and malignancies, a better understanding of the function of cilia in the processing and activation of Gli proteins is crucial.

HH signalling is essential for epidermal appendage formation in mammalian skin. Previous studies in Shh^-/-^ and Gli2^-/-^ mice revealed that Shh signalling, although dispensable for HF initiation, is required for HF down-growth past the hair germ stage of appendage formation. In particular, epithelial proliferation is decreased in developing HFs in both in Shh^-/-^ and Gli2^-/-^ mice (St-Jacques et al. 1998; Chiang et al. 1999, Mill et al. 2003). The expression of Gli1, one of the three transcription factors and also target gene of active HH signalling, is regulated by epithelial Gli2 activity in the skin. However, the loss of Gli1 does not disrupt mouse development, suggesting that Gli1 does not play a major role in mammalian skin development (Park et al. 2000; Bai et al. 2002; Mill et al. 2003). Earlier results implicated Gli2 as the primary activator downstream of HH signalling and key transcription factor mediating HH responses in the epidermis (Mill et al. 2003). However, due to potential functional redundancy of Gli2 and Gli3, the specific roles of the individual transcription factors in the mammalian skin epithelium have not been reported.

During HF morphogenesis, the HH response is induced in both, the epithelium and stromal cells, indicating that HH signal reception occurs in different HF compartments. Mouse models with a dermis-specific loss of Smo function showed the loss of dermal papilla precursor cells (Woo et al. 2012). Disrupting cilia assembly in the ventral dermis leads to an early arrest of HF development, similar to the HF defect observed in Shh-/- and Gli2-/- mice (Lehman et al. 2009; St-Jacques et al. 1998; Chiang et al. 1999, Mill et al. 2003). The *Ift88* and *Kif3a* cilia mutant mice with small or absent dermal condensates revealed that dermal cilia are critical signalling components for normal HF morphogenesis (Lehman et al. 2009). More recently it has been shown that HH activation in dermal papilla fibroblasts by expressing an active Smo mutant can induce new hair growth and increase fibroblasts heterogeneity (Liu et al. 2022). Genetic mouse experiments analysing IFT cilia mutants showed that the molecular trafficking machineries can be segregated according to their function in either ciliogenesis or in mediating HH signalling (Yang et al. 2015). However, how cilia function is linked to Gli processing and HH pathway activation in the mammalian skin epithelium still remains unclear.

Here, we use genetic approached to identify Gli2 as a crucial TF executing context-specific functions in skin appendage formation. Upon depletion of Gli2, but not Gli3, from the murine epidermis, HF morphogenesis is delayed whereas sebaceous gland (SG) formation is promoted, demonstrating a dual role for Gli2 in different HF sub-compartments. Mechanistically, our data reveal that Gli2 specifically regulates proliferation of SG stem cells, thereby controlling the development and maintenance of the number and size of individual SG lobes. Ablation of both, Gli2 and Gli3 in the skin epithelium, suggests redundant functions for both TFs. HF morphogenesis and differentiation are blocked in double knockout mice, mimicking the skin phenotype observed upon epidermal depletion of the HH receptor Smo. Finally, our work unravels cilia-independent Gli activator functions of Gli3 in Gli2 epidermal knockout mice.

## RESULTS

### Keratinocyte-specific ablation of Gli2, but not Gli3, causes defective hair follicle formation

To test the role of Gli transcription factors in the murine epidermis, we crossed Gli2^fl/fl^ mice with K14-Cre transgenic mice that express Cre recombinase from embryonic day (E) 12 onwards (Corrales at al. 2006; Hafner et al. 2004). At birth, K14Cre^tg^;Gli2^fl/fl^ mice (thereafter referred to as Gli2^EKO^) lacking Gli2 specifically in keratinocytes were macroscopically indistinguishable from Gli2^fl/fl^ littermates (insets Figure 1A). However, histological analysis of back skin tissue revealed that HF development was just initiated in Gli2^EKO^ new-born mice at postnatal day 0 (P0), whereas HFs were already growing deep into the underlying dermal tissue in control mice, indicating a strong delay or block in HF morphogenesis in Gli2^EKO^ animals (Figure 1A). The hair growth defect became macroscopically visible with impaired hair coat formation in Gli2^EKO^ mice at postnatal day 6 (P6) (Suppl Figure 1A). Analysis at P6 further demonstrated that the number of HFs was significantly reduced in Gli2^EKO^ mice (Figure 1B,C). Next, we investigated whether the lower HF number was due to a growth defect of a particular type of pelage hair. Compared to control littermates, Gli2^EKO^ mice showed a dramatic decrease in zigzag hairs and a slight increase in auchene/awl type of HF, while guard hairs were not changed (Figure 1D). Our data demonstrate that epidermal Gli2 activity is required for HFs forming in the second and third wave of pelage hair morphogenesis, starting at E16.5 and E18.5, respectively.

**Figure 1.**
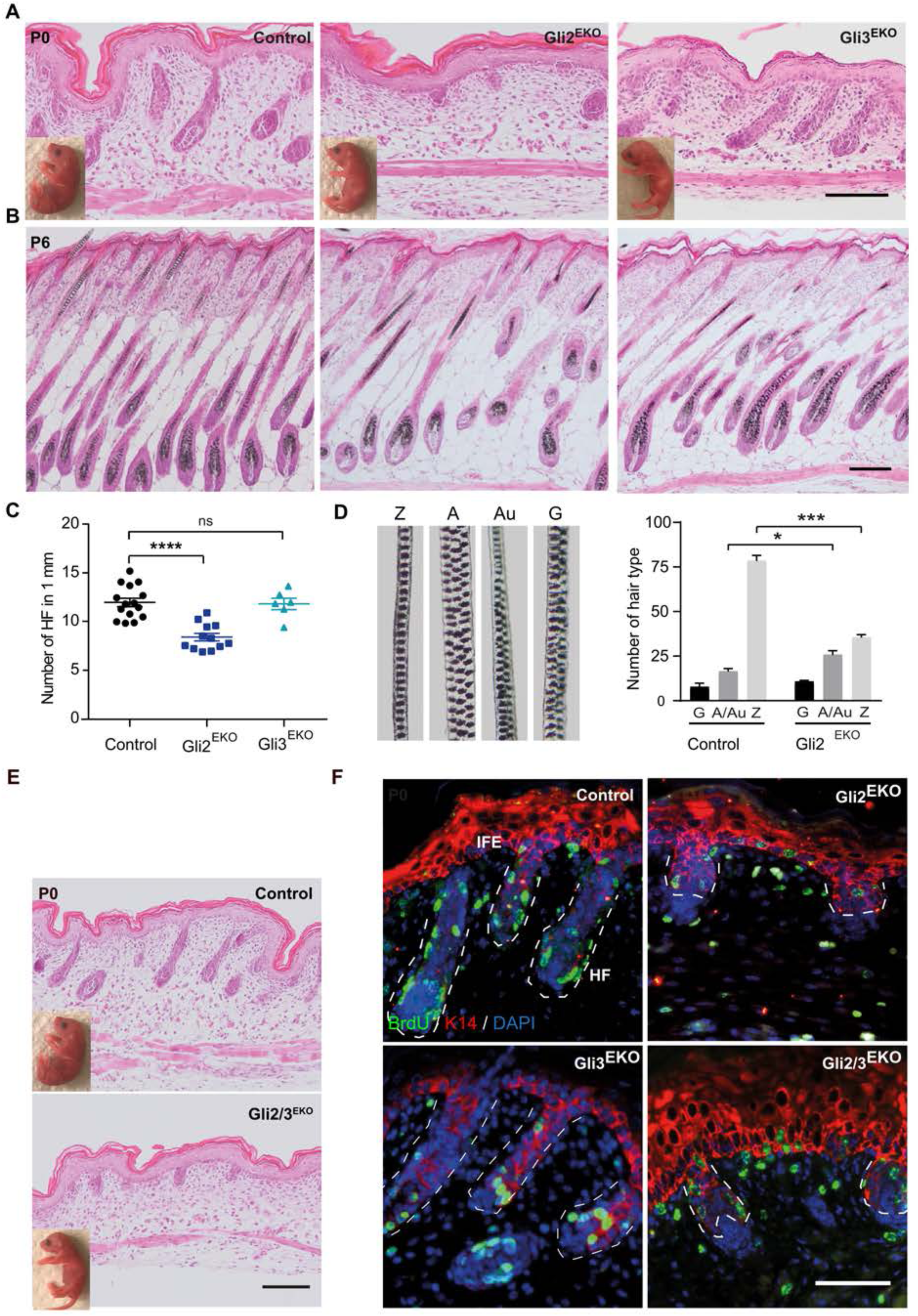
Synergistic functions of Gli2 and Gli3 are crucial for normal hair follicle morphogenesis. **(A,B)** Representative H&E staining of Gli2 (Gli2^EKO^) and Gli3 (Gli3^EKO^) epidermal knock out and control back skin at P0 **(A)** and P6 **(B)**. Inlets in (A): images of newborn animal at P0 (n = 3 mice/group), scale bar 100 µm. **(C)** Quantification of hair follicles in 1 mm back skin of Gli2^EKO^, Gli3^EKO^ and control mice (n = 3-6 mice/group). **(D)** Representative images of Zigzag (Z), Awl (A), Auchene (Au) and Guard (G) hair types in wild type skin and quantification of different hair types in P16 Gli2^EKO^ and control mice (n = 3 mice/group). **(E)** Representative H&E staining of Gli2/3^EKO^ and control littermates at P0. Inlets: images of P0 newborn mice (n = 3 mice/group). **(F)** Immunofluorescence staining for BrdU (green), K14 (red) and DAPI (blue, nuclei) of Gli2/3^EKO^ and control mice. (n = 3-4 mice/group), scale bar 50µm. *p<0.05, ***p< 0.001, ****p<0.0001. Data are presented as mean ± SEM.

To understand the role of epidermal Gli3 in appendage formation, we also generated a conditional epidermis-specific knockout for Gli3 (thereafter Gli3^EKO^ mice) (Blaess at al. 2008; Hafner et al. 2004). Gli3^EKO^ mice could not be distinguished from their littermate controls shortly after birth (insets in Figure 1A) and Gli3^EKO^ skin histology appeared similar to that of control animals. In contrast to Gli2^EKO^ mice, HF morphogenesis, including HF number, appeared normal in Gli3^EKO^ animals (Figure 1B,C), consistent with the phenotype seen in Gli3^-/-^ mice (Mill et al. 2003). Together the data indicate an important role for epidermal Gli2, but not Gli3 in the early stages of secondary and tertiary HF morphogenesis.

### Redundant functions of Gli2 and Gli3 in hair follicle morphogenesis

To address the possibility that Gli2 and Gli3 exert redundant functions and compensate for each other in the mouse epidermis, we generated Gli2/3^EKO^ double knockout mice by depleting both transcription factors from the epidermal tissue. Unlike Gli2^EKO^ and Gli3^EKO^ single mutants, Gli2/3^EKO^ double mutants had significantly deecreased body weight and showed severe skin abnormalities that became macroscopically apparent at P6 (Suppl Figure 1A-D). At birth, Gli2/3^EKO^ mice showed abnormal and blocked hair growth (Figure 1E). However, skin abnormalities became much more severe compared to control and Gli2^EKO^ mice (Figure 2A), where Gli2/3^EKO^ mice did not show any hair growth (Suppl Figure 1E). The skin defects were particularly prominent starting at P6, a time when control littermates had fully developed HFs reaching into the skin fat layer, and Gli2/3^EKO^ mice exhibited deformed epithelial invaginations and cyst-like structures that were not growing into the fat tissue of the skin (Suppl Figure 1E). These results demonstrate that epidermal Gli2 and Gli3 together are required for appendage formation in the mammalian skin. The data also point to redundant functions for Gli2 and Gli3 because single knockout mice showed milder (Gli2^EKO^ mice) or no (Gli3^EKO^ mice) skin phenotypes.

**Figure 2.**
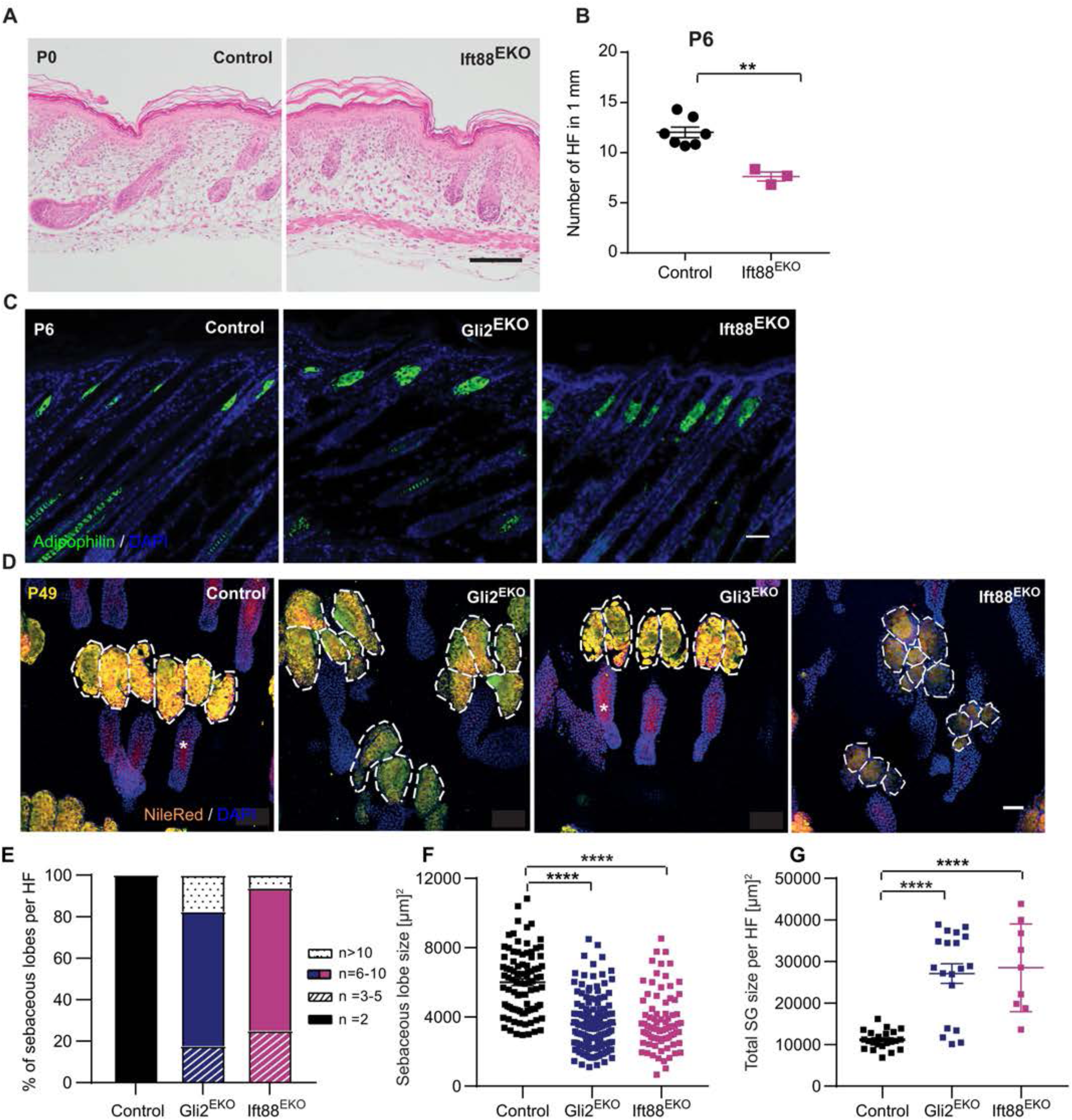
Cilia-dependent Gli2, but not Gli3 signalling in the skin appendage formation and maintenance. **(A)** Representative H&E staining of P0 back skin tissue from Ift88^EKO^ and control mice. (n = 3 mice/group)**. (B)** Quantification of hair follicles within 1 mm back skin of P6 Ift88^EKO^ and control mice (n = 3 mice/group). **(C)** Immunofluorescence staining for Adipophilin (green) and DAPI (blue, nuclei) of P6 back skin tissue from Ift88^EKO^ and control mice. **(D)** Nile red staining and DAPI (blue, nuclei) of P49 epidermal whole mounts from tail skin of Gli2^EKO^, Gli3^EKO^, Ift88^EKO^ and control mice. Dashed lines mark the individual lobes of sebaceous glands. (n = 3mice/group). **(E)** Quantification of number of individual sebaceous lobes per hair follicle of tail skin at P49. **(F)** Measurement of the size of individual sebaceous lobes and **(G)** total gland size in Gli2^EKO^, Ift88^EKO^ and control tail skin. Each dot represents an individual HF. Scale bars 100 µm (A), 50 µm (C,D). **p<0.01, ****p<0.0001. Data are presented as mean ± SEM.

Because HH signalling is associated with cell proliferation in the skin (St-Jacques et al. 1998; Chiang et al. 1999; Mill et al. 2003; Park et al. 2018), we next investigated whether changes in HF progenitor cell proliferation were underlying the HF defects in Gli2^EKO^ and Gli2/3^EKO^ mutant mice. Analysis of BrdU incorporation in new-born mice showed strong reduction of BrdU- positive cells in developing HFs of both, Gli2^EKO^ and Gli2/3^EKO^, but not Gli3^EKO^ mice when compared to control littermates (Figure 1F; Suppl. Figure 1F). The decrease in proliferation was also reflected by the differences in HF morphogenesis: whereas the majority of HF of Gli3^EKO^ and control mice where mainly in stage 6 of HF development, follicles of Gli2^EKO^ and Gli2/3^EKO^ mice were still at stage 4 (Suppl. Figure 1G) (Paus et al. 1999; Saxena et al. 2019).

The contribution of both, Gli2 and Gli3 to normal appendage formation in mammalian skin prompted us to investigate whether these two transcription factors are mediating the collective HH signalling response in the mammalian epidermis. We thus asked how the phenotypes of the individual and double Gli knockout mice compare to the skin phenotype of mice with the epidermal deletion of Smo, the membrane receptor for HH signalling (thereafter Smo^EKO^ mice). Smo^EKO^ mice were generated by crossing Smo^fl/fl^ line with the K14-Cre mouse line (Long et al. 2001; Hafner et al. 2004). Importantly, the skin phenotype of Smo^EKO^ new-born mice looked very similar to Gli2/3^EKO^ mice with an arrest in HF formation at the hair germ stage of morphogenesis (Suppl. Figure 1H). Our observations in Smo^EKO^ mice are supported by previous reports showing that depletion of Smo from mouse epidermis leads to defects in HF formation (Lichtenberger et al. 2016; Gritli-Linde et al. 2007).

### Gli2 and Gli3 are required for hair lineage differentiation and hair follicle stem cells

Next, we investigated whether Gli transcription factors play a role in HF stem and progenitor lineage differentiation and analysed the expression of specific hair differentiation markers at P6, when HF progenitor differentiation takes place. Immune histological analysis detected keratin (K) 75, a marker for the HF companion layer, in control as well as Gli2^EKO^ and Gli3^EKO^ mice. Further, the HF inner root sheath marker K71 and the hair cortex marker K86 were normally expressed in Gli2^EKO^ and Gli3^EKO^ and littermate control mice (Suppl Figure 2A), demonstrating that hair lineages were normally generated in Gli2^EKO^ and Gli3^EKO^ mice. RNA analysis showed that the expression of K75, K71 and K86 were significantly reduced in Gli2^EKO^ mice compared to control and Gli3^EKO^ animals (Suppl Figure 2B,C), most likely due to the delayed HF morphogenesis and reduction in HF number observed in Gli2^EKO^ mice (Figure 1A-C).

In contrast, expression of hair lineage markers was completely blocked in Gli2/3^EKO^ mice when compared to control skin samples (Suppl Figure 3A). A block in hair differentiation marker expression was also detected at the mRNA level (Suppl Figure 3B) indicating that HF progenitor cells do not undergo hair lineage differentiation in Gli2/3^EKO^ mice. We then asked what epithelial cell types populate the abnormal HFs and cyst-like structures observed in Gli2/3^EKO^ mutants. Interestingly, K10, a marker for suprabasal keratinocytes within the interfollicular epidermis, was detected in cyst-like and abnormal HF structures in Gli2/3^EKO^ mice, but not in HFs of littermate controls (Suppl Figure 3C). Expression of interfollicular markers K10 and filaggrin was also strongly elevated at the RNA level (Suppl Figure 3D,E). These results indicate that both transcription factors, Gli2 and Gli3 together are required for HF progenitor lineage differentiation. Given that hair differentiation was completely blocked in Gli2/3^EKO^ mice, we wondered whether HF SCs are generated in Gli2/3^EKO^ mice. To address this question, HF bulge SC marker K15 was analysed in adult mice. As expected, K15 was strongly expressed by bulge SCs localising to the lower permanent part of the HF in control animals. However, K15 protein was not detected in abnormal cyst-like epithelial structures in Gli2/3^EKO^ mice (Suppl Figure 3F), demonstrating that Gli2 and Gli3 together are critical for the establishment of bulge SCs and progenitor differentiation in distinct HF cell types. This conclusion was further supported by qRT-PCR experiments showing reduced mRNA expression for K15 (Suppl Figure 3G), whereas transcription of HF bulge marker SOX9 was not significantly changed in Gli2/3^EKO^ skin (Figure 3H).

**Figure 3.**
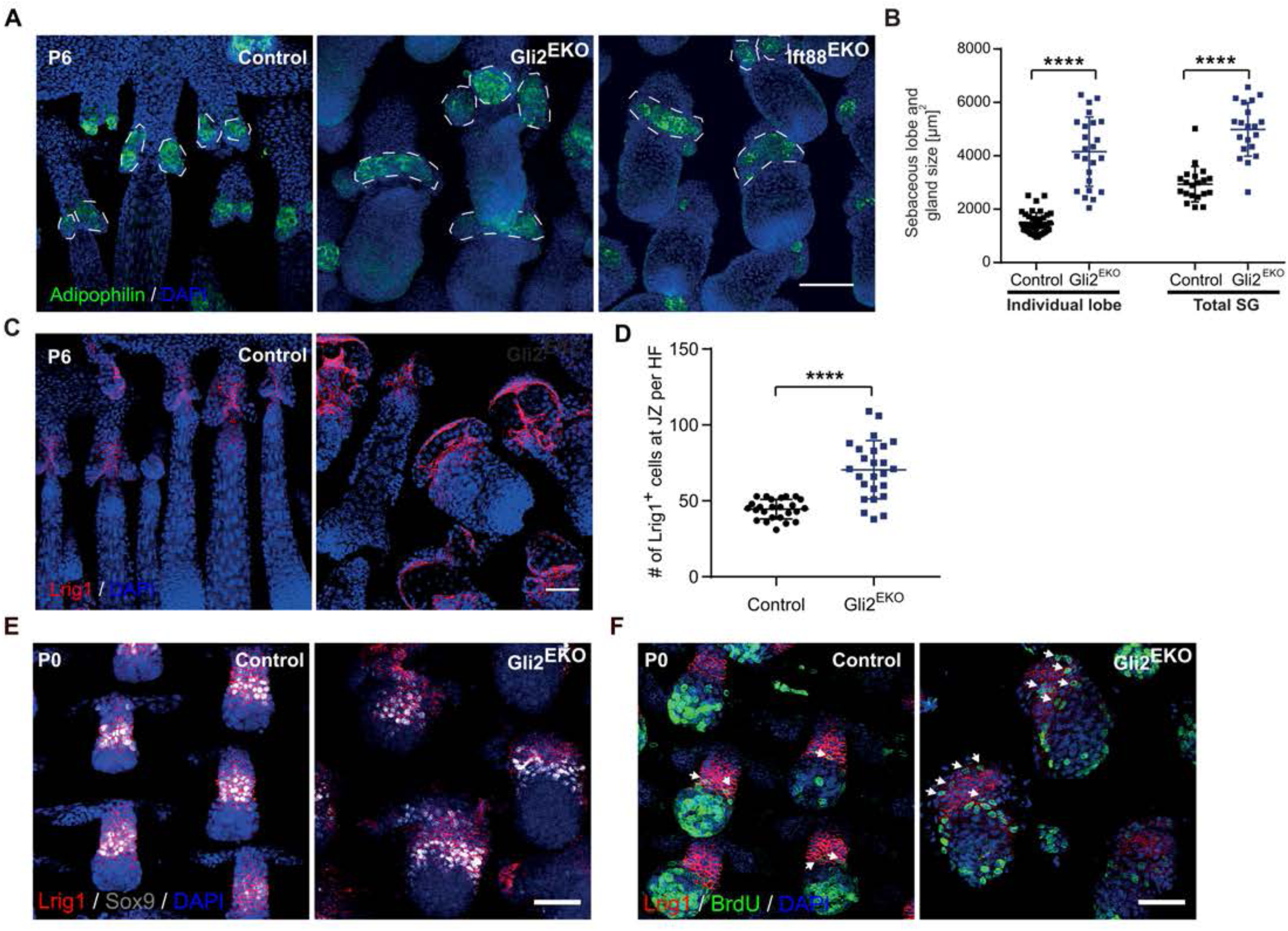
Gli2 regulates sebaceous gland stem cells during appendage formation. **(A)** Immunofluorescence staining for Adipophilin (green) and DAPI (blue, nuclei) of epidermal whole mounts of tail skin of P6 Gli2^EKO^, Ift88^EKO^ and control tail skin. Dashed lines mark the developing SGs in tail HFs. **(B)** Quantifications of the size of the individual sebaceous lobes (left) and total sebaceous gland (SG) (right) per hair follicle at P6. **(C)** Immunofluorescence staining for Lrig1 (red) and DAPI (blue, nuclei) of epidermal whole mounts of tail skin of P6 Gli2^EKO^ and control mice (n = 3-4 mice/group). **(D)** Quantification of the Lrig1^+^ cells per hair follicle. **(E,F)** Immunofluorescence staining of epidermal whole mounts for Lrig1 (red), Sox9 (grey) **(E)** and BrdU (green), Lrig1 (red) and DAPI (blue, nuclei) **(F)** of Gli2^EKO^ and control tail skin. Scale bar 50 µm (A,C,E,F). ****p<0.0001. Data are presented as mean ± SEM.

### Cilia-dependent Gli2, but not Gli3 signalling in the skin epithelium

Given that cilia play an important role in conducting the HH signal (Lee et al. 2016; Zhang and Beachy 2023), we asked which role do cilia play in mediating epithelial Gli transcription factor signalling in skin appendage formation? A cilia mutant Ift88^EKO^ mouse line was analysed, depleting Ift88 from the skin epithelium by crossing Ift88^fl/fl^ with K14Cre animals (Haycraft et al. 2007; Hafner et al. 2004). Importantly, Ift88^EKO^ mice do not form cilia in the skin epithelium (Damen et al. 2021; Croyle et al. 2011). Initial HF formation was delayed in Ift88^EKO^ new-born mice compared to control littermates (Figure 2A). Later at P6, HF developed further and looked similar to those of control animals. However, the number of HF were significantly reduced in Ift88^EKO^ mice (Figure 2B). The phenotype seen in new-born and young Ift88^EKO^ mice appeared highly similar to the HF phenotype observed in Gli2^EKO^ animals (Figure 1A-C).

### Specific role for epidermal cilia-Gli2 signalling axis in the formation and maintenance of sebaceous glands

The resemblance of the skin phenotype seen in Gli2^EKO^ and Ift88^EKO^ mice also extended to the formation of sebaceous glands (SG), lipid producing skin appendages normally attached to the HFs. At P6, SGs visualised by adipophilin staining appeared larger in the back skin of both Gli2^EKO^ and Ift88^EKO^ mice when compared to the littermate controls (Figure 2C). These data show that in contrast to HF development, the formation of SGs was not delayed but rather accelerated in Gli2- and cilia-defective mice pointing to rather a cell-type specific and context-dependent effect of HH signalling.

We wondered whether this is a transient defect during mouse development or whether Gli2 signalling plays a general role in SG renewal and maintenance? To address this issue, we examined epidermal whole mounts of tail skin in adult mice, where two prominent sebaceous glands are attached to one HF and are easily accessible for a more detailed characterisation. Our analyses revealed that the Nile Red-positive SGs were clearly enlarged in 7 weeks old Gli2^EKO^ mice when compared to control littermates (Figure 2D). In contrast, Gli3^EKO^ mice displayed SGs of normal size (Figure 2D). Next, we wanted to know whether SGs were also affected in adult cilia mutant mice. Interestingly, SGs of adult Ift88^EKO^ mice were also enlarged and resembled the phenotype observed in Gli2^EKO^ mice of the same age (Figure 2D).

Quantitative analysis revealed that the majority of SGs attached to one HF in Gli2^EKO^ and Ift88^EKO^ mice consisted of 6 to 10 individual lobes, whereas glands in control littermates always contained 2 lobes. Some HFs in both mutant mouse lines contained SGs made of 10 or more lobes (Figure 2E). The individual SG lobes were significantly smaller in Gli2^EKO^ and Ift88^EKO^ mice (Figure 2F). However, due to the dramatic increase in the number of lobes, the overall volume of the SGs attached to one HF was evidently increased in Gli2^EKO^ and Ift88^EKO^ mice when compared to littermate controls (Figure 2G). Together, the similar phenotype seen in Gli2^EKO^ and Ift88^EKO^ mice points to cilia-dependent signalling of Gli2 in SG morphogenesis and maintenance (Croyle et al. 2011).

### Gli2 signalling regulates sebaceous gland stem cells during appendage formation

Next, we investigated the potential role of Gli2 during SG morphogenesis using epidermal tail whole mounts. We focussed on P6, when SG formation is initiated and the first mature sebocytes emerge within the upper part of the HF (Frances and Niemann, 2012). Staining for the sebocyte marker adipophilin revealed that the number of developing lobes were increased in Gli2^EKO^ and Ift88^EKO^ mice already during appendage formation (Figure 3A), supporting our finding at P49 (Figure 2D). The developing lobes localised to the same region of the developing HF as seen in control mice. Quantifications showed that the size of the individual lobes and the entire SGs were significantly increased to more than twice as big in Gli2^EKO^ and IFT88^EKO^ mice compared to littermate controls (Figure 3B).

Furthermore, we wanted to find out whether Gli2 signalling controls the SC compartment that gives rise to mature sebocytes and SGs. Sebocytes have been previously shown to originate from Lrig1+ve keratinocytes during SG morphogenesis, we thus analysed epidermal whole mounts for the expression of Lrig1 (Frances and Niemann, 2012; Andersen et al. 2019). As expected, Lrig1 was strongly expressed in the junctional zone, adjacent to and surrounding the base of the two sebocyte clusters forming in the upper part of developing HFs at P6 control mice (Figure 3C). In contrast, the Lrig1+ve cell population was dramatically increased and expanded within the upper region of HFs, perhaps giving rise to the multiple lobes in the SGs of Gli2^EKO^ mice (Figure 3C). Quantitative analysis revealed a significant increase in the number of Lrig1+ve cells localising to the junctional zone of the HF at P6 in Gli2^EKO^ mice when compared to control animals (Figure 3D). These data demonstrate that epithelial Gli signalling is required for normal SG development by restricting the size and localisation of the Lrig1+ve stem cell compartment during skin appendage formation.

Next, we addressed whether the establishment of different HF SC compartments is affected in epidermal Gli2^EKO^ mice. Therefore, we analysed HF morphogenesis (P0) during the transition of early epithelial HF progenitors expressing the markers SOX9 and Lrig1 into distinct SC compartments of the future HF bulge (SOX9) and the junctional zone (Lrig1) at P0 of development (Frances and Niemann 2012). The process of SOX9+ve cells separating from Lrig1+ve SG progenitors appeared similar in control and Gli2^EKO^ mice (Figure 3E). Further, we investigated which SC compartment undergoes cell proliferation in the absence of epidermal Gli2. BrdU incorporation experiments revealed that proliferation takes place in HF cells below the Lrig1+ve compartment in Gli2^EKO^ and littermate controls. Remarkably, a dramatic increase in the number of BrdU+ve cells was detected within the Lrig1 compartment in Gli2^EKO^ skin samples when compared to control mice (Figure 3F, arrows), indicating that Gli2 is regulating SG formation specifically by controlling the proliferation of Lrig1+ve SCs.

### Gli3 activator function in epidermal appendage formation

We finally wanted to gain more mechanistic insights into the redundant functions of Gli2 and Gli3 in the skin epidermis and aimed to better understand how Gli3 exerts compensatory activities in Gli2^EKO^ mice. To address this, we tested the posttranslational processing of full-length Gli3 protein into the Gli3 repressor (Gli3^R^) (Zhang and Beachy, 2023). Western blot analysis revealed a strong Gli3^R^ band (83 kDa) that was absent in epidermal samples from Gli3^EKO^ embryos (suppl. Figure 4A,B), demonstrating the specificity of the antibody and detected band. Interestingly, Gli3^R^ protein was strongly reduced in the isolated epidermis from Gli2^EKO^ embryos compared to control littermates indicating that Gli3 might function as transcriptional activator in Gli2^EKO^ mice (Figure 4A; Suppl. Figure 4A,C). This was further strengthened by the analysis of the HH target gene Gli1. Whereas Gli1 mRNA levels did not significantly differ in Gli2^EKO^ and Gli3^EKO^ mice compared to control littermates, Gli1 expression was strongly reduced in Gli2/3EKO mice. These data show that HH signalling was still active in Gli2^EKO^ mice and demonstrate that only the depletion of Gli2 and Gli3 together results in block of HH signalling in the skin epithelium (Figure 4C).

**Figure 4.**
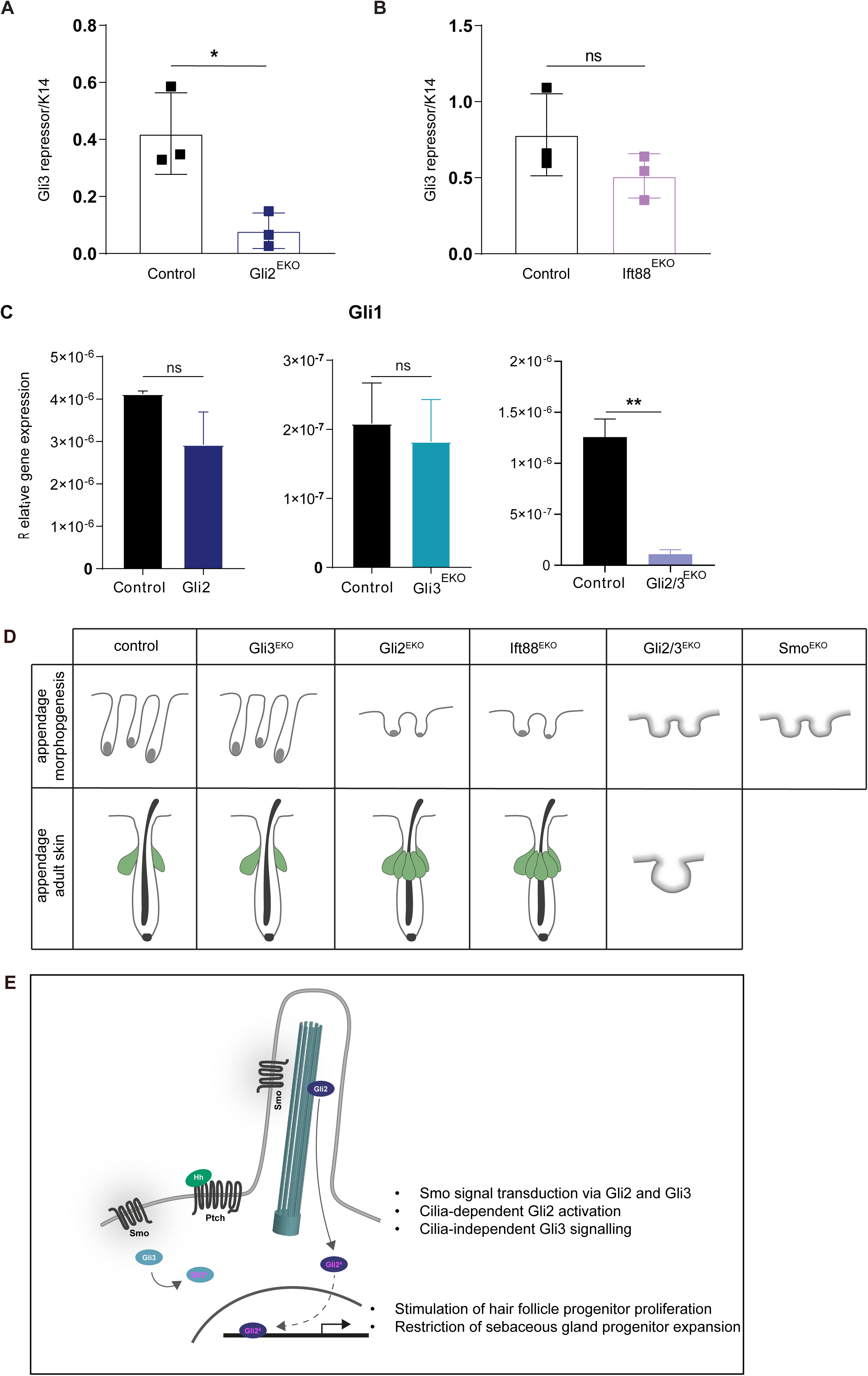
Gli3 activator function in epidermal appendage formation. **(A,B)** Quantification of Gli3^R^ in epidermis of Gli2^EKO^ **(A)** and Ift88^EKO^ mice **(B)** based on western blot analysis (shown in Suppl Fig 4) and normalized to detection of K14 protein. *p<0.05. Data are presented as mean ± SD. **(C)** Quantification of Gli1 target gene expression in different Gli mouse mutants (n=?). **p<0.01. Date are presented as mean ± SD. **(D)** Schematic overview dissecting phenotypic characterization during skin appendage development and adult skin of different Hh signaling mouse mutants, including Gli2^EKO^, Gli3^EKO^, Ift88^EKO^, Gli2/3^EKO^ and Smo^EKO^ mice. **(E)** Schematic summary of distinct Gli2 signaling activities in mammalian appendage formation and maintenance.

Given our previous observation that the defects in epidermal appendage formation are very similar in Gli2^EKO^ and Ift88^EKO^ mice, we wondered whether Gli3 protein processing is also affected in Ift88^EKO^ mice? The Gli3 western blot showed no statistically significant reduction in the Gli3^R^ form (Figure 4B, Suppl. Figure 4C,D). Our data suggest that the reduction in Gli3^R^ in Gli2^EKO^ might alleviate the HH phenotype in these mutants compared to Gli2/3^EKO^ mice.

## DISCUSSION

A question of fundamental importance is how epithelial appendages are generated and organise into different cell types and functional tissue compartments (Abe and Tanaka 2017). Using a variety of different genetic knock out mouse models, this study dissects HH-driven molecular and cellular mechanisms underlying epidermal appendage formation. To our knowledge, our study shows for the first time a dual and context-dependent role for the transcription factor Gli2 during the morphogenesis of the pilo-sebaceous unit in the mammalian skin epithelium. On one hand, epidermal Gli2 is required for the proliferation of HF progenitors, whereas on the other hand, Gli2 restricts the expansion of SG progenitor cells. Based on our observation that early defects in HF formation seen in Gli2^EKO^ and Gli2/3^EKO^ new-born mice appeared very similar, the data suggest that Gli2 drives progenitor proliferation in HF keratinocytes at the early stages of HF morphogenesis. Furthermore, our results reveal that the initial block in HF formation in Gli2^EKO^ mice is likely rescued by Gli3 at later stages of morphogenesis, and demonstrating compensatory functions in the skin epithelium. Thus, our work identifies an important role for epidermal Gli3 in skin appendage formation, a HH transcription factor that has previously been neglected in skin research, in the absence of Gli2 in skin appendage formation.

Our comprehensive analysis of genetic mouse models revealed a similar phenotype of defective appendage formation in Gli2/3^EKO^ and Smo^EKO^ mice and thus uncovered that synergistic signalling by Gli2 and Gli3 is predominantly mediating HH signalling in mammalian skin epithelium. Our results also support the previous notion that Gli1 does not seem to play a major role in transducing a HH response in mammalian epidermis and thus is not contributing to appendage formation in mammalian skin.

Our work identifying an essential role for Gli2 in controlling the number and localisation of Lrig1+ve stem cells driving SG morphogenesis is of particular interest given that previous data pointed to a role for HH signalling in governing SG differentiation in adult skin. First evidence supporting this hypothesis came from Shh mutant mice that showed deficiency in SG formation (St. Jacques et al. 1998; Chiang et al. 1999). This observation was further supported by a mouse model with epidermis-specific overexpression of an active Smo mutant receptor resulting in the increase in size and number of SGs in adult skin (Allen et al. 2003). In contrast, blocking HH signalling in epithelial skin cells by overexpressing a Gli2 mutant suppresses sebocyte differentiation (Allen et al. 2003). Interestingly, expression of an active Gli2 mutant in the skin epithelium induces a SG duct cell fate and branched and additional SGs in adult mice (Gu and Coulombe, 2008). Furthermore, a role for HH-Gli in differentiation was also detected in a human sebocyte cell line and SG tumours of human and mice (Niemann et al. 2003; Kakanj et al. 2013; Takeda et al. 2006). Together these studies demonstrated that modulating the level of HH signalling affects SG physiology, how this is mechanistically regulated was not known. Our novel results suggest that Gli2 could exert differential functions during SG morphogenesis and SG maintenance in adult skin: whereas Gli2 restricts the expansion of SG stem cells during appendage formation, Gli2 and HH activation promotes SG differentiation in adult skin and skin tumours (Kakanj et al. 2013). Remarkably, the SG defect observed in Gli2^EKO^ mice is maintained throughout adult live and multiple cycles of hair regeneration showing that early Gli2-mediated regulation of SG formation is relevant for adult skin physiology.

Our observation that Gli2/3^EKO^ mice showed a more severe phenotype when compared to the Gli2 single knockout mouse model strongly suggests a compensatory role for the Gli3 transcription factors mediating the HH response during appendage formation. Moreover, the lowering of the Gli3^R^ in Gli2^EKO^ mice points to an under-appreciated Gli3 activator function compensating for Gli2 in HF formation and growth. This hypothesis was further supported by Gli1 target gene expression that is only blocked in Gli2/3^EKO^ mice, demonstrating that only the depletion of Gli2 and Gli3 together from the skin epithelium results in block of HH signalling.

Given prior reports showing HH responsiveness particularly in the dermis, our work dissects and highlights its importance in the skin epithelium. We demonstrate that Gli2 functions, but not Gli3, are cilia-dependent in the process of appendage formation. Intriguing in this respect is that the Ift88 dermal knockout recapitulates the HF growth defect seen in Shh and Gli2 complete knockout mice (Lehman et al. 2009), whereas the Ift88^EKO^ mutants phenocopy the milder hair phenotype also observed in Gli2^EKO^ animals. Together these data reveal that either Gli3 does not play a role in the dermal compartment or that cilia play differential roles in transducing the HH signal and are more important in dermal cells. A context-dependent role of primary cilia transducing the HH signal has been proposed previously where cilia can act as both positive and negative regulators of the HH signalling pathway. In the neural tube, where Gli activators normally play an important role, defects in cilia and IFT result in loss of function HH phenotypes, whereas in the limb, where Gli3 repressors have a major role, these defects lead to a gain of function HH phenotype (Haycraft et al. 2005; Huangfu and Anderson, 2005). Our results suggest that HF and SG development are differentially regulated by the cilia-Gli2 signalling axis and indicate that Gli2 signalling is mediated by cilia. Moreover, given that the skin phenotype is much more severe in Gli2/3^EKO^ mice when compared to Ift88^EKO^ animals, the results also suggest that Gli3 functions independent from cilia in the mammalian skin epithelium. Our results showing that cilia do not play an import role in processing full-length Gli3 into the Gli3^R^, further support cilia-independent Gli3 functions in skin appendage formation.

## Acknowledgements

We are grateful to Sandra Blaess (University of Bonn) for providing floxed Gli mouse lines and Beate Lichtenberger (University of Vienna) for the floxed smoothened mouse line.

We thank Houda Khatif for excellent experimental support and Deborah Delbue to assist in manuscript preparation. We thank for the support by the animal facilities of CECAD, CMMC and the Medical Faculty and the CMMC/CECAD Imaging Facility.

Work of H. Bazzi was funded by the German Research Foundation (SFB829, Project-ID 73111208). Research of C. Niemann was funded by the LEO Foundation (LF-OC-22-001112) and the German Research Foundation (SFB829, Project-ID 73111208 and DFG grant, Project-ID 507956072). C Niemann is grateful for financial support by the Köln Fortune Program of the Medical Faculty (Project-ID 342/2019) and the CMMC of the University of Cologne.

## Material and Methods

### Experimental mice

Epidermal deletion of Gli2, Gli3, and Smo and Ift88 was accomplished by crossing K14Cre^tg^ mouse line (Hafner et al. 2004) with Gli2^flox^ (Corrales et al. 2006), Gli3^flox^ (Blaess et al. 2008), Smo^flox^ (Long et al. 2001), and Ift88^flox^ (Haycraft et al. 2007) lines which have been described previously. All mouse strains except Smo^flox^ and Ift88^EKO^ were maintained and crossed on the C57BI/6N background. Ift88^EKO^ mice were maintained and crossed on mixed FVB/N and C57Bl/6N background. Floxed littermates without Cre recombinase were used as a control without gender preference. All experiments were conducted with samples from three independent litters. All husbandry and animal experiments were conducted according to guidelines and license approval by the State Office North-Rhine Westphalia, Germany.

### BrdU labeling

A single dosage of BrdU (Sigma; 100 mg/kg body weight) was administered i.p. one hour prior sacrificing the animals. BrdU+ve cells were counted together with the total number of cells (DAPI+ve) and their ratio was calculated as the percentage of BrdU+ve cells.

### Histological analysis

Harvested back skin samples were fixed in 4% formaldehyde solution for 2 hours. Paraffin-embedded samples were sectioned with 5 µm thickness, and hematoxylin & eosin (H&E) staining was performed. Images were taken with BX53 microscope (Olympus) and analyzed with ImageJ software.

H&E images of P6 back skin samples (approximately 1cm) were taken with BX53 microscope (Olympus) and analyzed with ImageJ software. HFs with attached SG or dermal papillae were counted from 3 different fields on each skin sample. HF numbers were normalized to covered skin area and presented as means of different measurements.

For hair type analysis, hairs from three different areas along the central axis were plugged and mounted on slides. Hair types were determined based on the patterns and bends of the hair shaft. For each genotype, at least three mice were analyzed.

### Isolation of tail epidermal whole mount

As described previously (Braun et al. 2003), tail skin was incubated with 5mM EDTA for 1h (P6 animals) and 3h (for older animals) at 37^°^C. The tail epidermis was removed carefully with a tweezer and fixed in 3.4% formaldehyde solution for 1h. Tail epidermis samples were stored in PBS at 4°C for further analysis.

### Immunofluorescence staining

For immunofluorescence, paraffin tissue sections were incubated with 10% normal goat/donkey serum for 1h. Only for K71, K75 and K86 staining, 0.1% Triton X was included in blocking solution. Tail epidermal whole mount were incubated with TB Buffer (0.25% Fish Skin Gelatin, 0.005 Triton X, 0.25% milk powder in TBS) for 1h. The following primary antibodies were used: K14 (rabbit, 1:1000; Covance), K10 (rabbit, 1:1000; Covance), K71 (g.pig, 1:200, Progen), K75 (g.pig, 1:200, Progen), K86 (g.pig, 1:200, Progen), K15 (g.pig, 1:1500, Progen), Sox9 (rabbit, 1:1000, Abcam), Adipophilin (g,pig, 1:250, Fitzgerald Industries) BrdU (mouse, 1:20, BD), Lrig1 (goat, 1:100, R&D). All secondary antibodies were coupled either Alexa-488 or Alexa-594 (1:500, Invitrogen). 1:1000 DAPI (20mg/ml, Sigma-Aldrich) was used to counterstain the nuclei.

Nile red solution (1mg/ml in acetone) was freshly diluted 1:1000 in PBS and tail epidermal whole mounts were incubated for 30 minutes at room temperature before rinsing a few times with PBS. For nuclei counterstaining, DAPI was used.

All images were taken with IX83 or FV1000 (Olympus) and analyzed with ImageJ software and Adobe Illustrator CS5.

### Quantification of sebaceous gland size and number

Following Nile red staining, images were taken randomized of at least 3 different fields from the tail epidermis with FV1000 (Olympus). The individual size of the sebaceous lobes was measure with ImageJ software focusing on the Nile red/Adipophilin positive area. SGs with low integrity and no clear separation were excluded from the analysis. Based on the SG staining, the number of individual glands per HF were also calculated.

### qRT-PCR analysis

RNA isolation was done with RNAMagic (Biobudget), and cDNA was synthesized from 0.5 µg total RNA using Quantitech reverse transcription kit (Qiagen) according to the manufacturer’s protocol. All primers used for qRT-PCR were listed in Supplementary Table 1. Two sets of independent experiments with experimental triplicates were run on Quantstudio3 (Applied Biosystems). Relative gene expression was normalized to 18S housekeeping gene expression.

### Protein analysis by western blot

Embryonic back skin was dissected from mice at embryonic day E16.5. To separate epidermis and dermis, the skin was incubated for 30 min in 400 μl ice-cold split buffer containing 3.8% ammonium thiocyanate. To prepare lysates for WB, epidermal sheets were washed in 1x PBS and added to 50µl 2x Laemmli (BioRad #1610737) buffer before dissociating the tissue by pipetting and freezing on dry ice. Lysates were supplemented with 10% ß-mercaptoethanol and boiled for 5 min at 95°C before loading 10 µl per sample on 4-15% gradient gels (BioRad). SDS page was performed for 1h at 100V. Proteins were transferred onto PVDF membranes (0.45 µm pore size, GE Healthcare) for 16 h at 35 V. Membranes were stained for 5 min with Ponceau S (Sigma) before blocking with 5% milk in TBST for 1h at RT. Membranes were labeled with primary anti-Gli-3 antibody (1:1000, R&D) overnight at 4°C, followed by incubation of secondary HRP-conjugated anti goat antibody for 45 min at RT (1:10.000, Jackson ImmunoResearch). For detection of K14, membranes were incubated with anti-K14 (1:10.000, rabbit, BioLegend) overnight at 4°C, before incubating with anti-rabbit pig HRP (1:10.000, Cytiva) antibody for 45 min at RT. Detection of signal was performed with Ammersham Prime ECL (Cytiva).

Quantification of band intensities was performed with FiJi.

### Statistical analysis

Statistical significance was calculated using unpaired two-tailed Student’s t-test. *p < 0.05, **p < 0.01, ***p < 0.001 and ****p < 0.0001 were considered as statistically significant. All data were described as mean ± SEM or ± SD as indicated. All statistical analyses were done using GraphPad Prism 7 (GraphPad Software, Inc., La Jolla, CA, USA).

