## Supplemental table 1 and supplemental figures 1-4 for "Specific and redundant roles for Gli2 and Gli3 in establishing cell fate during hair follicle development"

**Supplementary Table Gozum et al.**
List of mouse primers for quantitative real-time PCR (qRT-PCR)

| Target | Primer | Sequence |
|---|---|---|
| 18S | Forward | 5`- ATCAGATACCGTCGTAG -3` |
| | Reverse | 5`- GCAAAGCTGAAACTTAAAG - 3` |
| Filaggrin | Forward | 5`- GACAGCCAAGTCCATTCT -3` |
| | Reverse | 5`- ACTCATTCCTCCCTGAC – 3` |
| Gli1 | Forward | 5'-GCACCACATCAACAGTGAGC-3` |
| | Reverse | 5'-GACTTCCGACAGCCTTCAAA-3` |
| Gli2 | Forward | 5`- CGCATGATTCGGACCTCTC -3 |
| | Reverse | 5`- GATTGATGGGGTGGGGAAAA -3` |
| Gli3 | Forward | 5`- AACTCCTTGGTTACAATCCTCAAT – 3` |
| | Reverse | 5`- GATAGGTCTCTGTGTTGGAAATGT – 3` |
| K10 | Forward | 5`- ACCCTTAGCAAGTCTGACCT - 3 |
| | Reverse | 5`- CGTTCATTTCCACATTCACATCAC - 3 |
| K71 | Forward | 5`- AGACTCCGCTCAGAGATTG- 3 |
| | Reverse | 5`- CATCCTTGAGGGCACTGT – 3` |
| K75 | Forward | 5`- CGCAACACCAAACAAGAG – 3` |
| | Reverse | 5`- GGCATCCTTGAGAGCTAG – 3` |
| K86 | Forward | 5`- GGAGCAGAGGTTGTGTGAGG - 3` |
| | Reverse | 5`- AGGGGCAGTACCAGAGACG – 3` |
| K15 | Forward | 5’- TGGAGATGCAGATTGAGCAGCTGAA - 3` |
| | Reverse | 5’ – TGCTCCCTCATCTCTGCCAGCA – 3` |
| Sox9 | Forward | 5`- CCTTCATGGTGTGGGCGCAG - 3` |
| | Reverse | 5` - GGGTGGTCTTTCTTGTGCTGCA - 3` |

**Supplementary Figures Gozum et al.**

**Supplementary Figure 1. Generation and characterization of genetic Gli and Smo mouse models.**

**(A,B)** Representative images **(A)** and body weight **(B)** of Gli2$^{EKO}$, Gli3$^{EKO}$, Gli2/3$^{EKO}$ and control mice at P6.

**(C,D)** Expression of Gli2 and Gli3 in mouse epidermis were depleted with Cre recombinase under control of K14 promoter. qRT-PCR analysis showing Gli2 **(C)** and Gli3 **(D)** mRNA expression in P6 Gli2/3$^{EKO}$ and control mice tail epidermis.

**(E)** Representative H&E staining of Gli2/3$^{EKO}$ and control littermates at P2 and P6.

**(F)** Quantification of BrdU$^{+}$ proliferative cells per DAPI$^{+}$ HF keratinocytes (n= 12-27 HFs/group).

**(G)** Quantification of HF stages in Gli2/3$^{EKO}$ and control mice at P0. (n=22-36 HFs/group).

**(H)** Representative H&E staining of back skin tissue from Smo$^{EKO}$ and control mice at P0.

Scale bar, 50µm (E), 100 µm (H).*p < 0.05, **p<0.01, ***p<0.001, ****p<0.0001, t-test, (n = 3 mice/group). Data are presented as mean ± SEM.

**Supplementary Figure 2. Analysis of hair lineage differentiation in Gli2$^{EKO}$ and Gli3$^{EKO}$ mice.**

**(A)** Immunofluorescence staining for K14 (green), K75 (red), K71 (grey), K86 (green) and DAPI (blue, nuclei) of P6 back skin tissue from Gli2$^{EKO}$, Gli3$^{EKO}$ and control mice. (n= 3 mice/group), scale bar 50 µm.

**(B,C)** qRT-PCR analysis of K75 , K71 and K86 mRNA expression in back skin samples Gli2$^{EKO}$ **(B)** and Gli3$^{EKO}$ **(C)** and control littermate at P6. *p<0.05, **P < 0.01, t-test, (n = 3 mice/group). Data are represented as mean ± SEM.

**Supplementary Figure 3. Gli2 and Gli3 are required for hair lineage differentiation and hair follicle stem cells.**

**(A)** Immunofluorescence staining for K14 (green), K75 (red), K71 (grey), K86 (green) and DAPI (blue, nuclei) of P6 back skin tissue from Gli2/3$^{EKO}$ and control mice. Dashed lines mark the hair follicle-like structures in Gli2/3$^{EKO}$ skin (n= 3 mice/group).

**(B)** qRT-PCR analysis for K75, K71 and K86 mRNA expression in back skin of Gli2/3$^{EKO}$ and control mice. (n = 3 mice/group).

**(C)** Immunofluorescence staining for K10 (red) and DAPI (blue, nuclei) of P6 Gli2/3$^{EKO}$ and control back skin.

**(D,E)** qRT-PCR analysis of K10 and Filaggrin mRNA expression in back skin of Gli2/3$^{EKO}$ and control mice. (n =3 mice/group).

**(F)** Immunofluorescence staining for K15 (red) and DAPI (blue, nuclei) of P49 back skin tissue from Gli2/3$^{EKO}$ and control mice

**(G,H)** qRT-PCR analysis of bulge marker K15 **(G)** and Sox mRNA expression **(H)** in back skin of Gli2/3$^{EKO}$ and control mice. (n =3 mice/group).

Scale bar, 50 µm (A,C,F). *p<0.05 **p < 0.01, ****p<0.0001. Data are represented as mean ± SEM.

**Supplementary Figure 4. Western blot images**

Representative western blot images of Gli3 **(A,C)** and K14 protein **(B,D)** analysis in epidermal lysates from Gli2$^{EKO}$ **(A,B)** and Ift88$^{EKO}$ **(C,D)** mice. Lysates of three independent control and mutant mice are presented. Lysates of control whisker pad epidermis and Gli3$^{EKO}$ were used as positive and negative control respectively for $Gli3^R$ detection. **(A,C)** Gli3R is detected at 83 kDa, unspecific bands are indicated by asterisks. **(B,D)** Detection of K14 at 53 kDa.

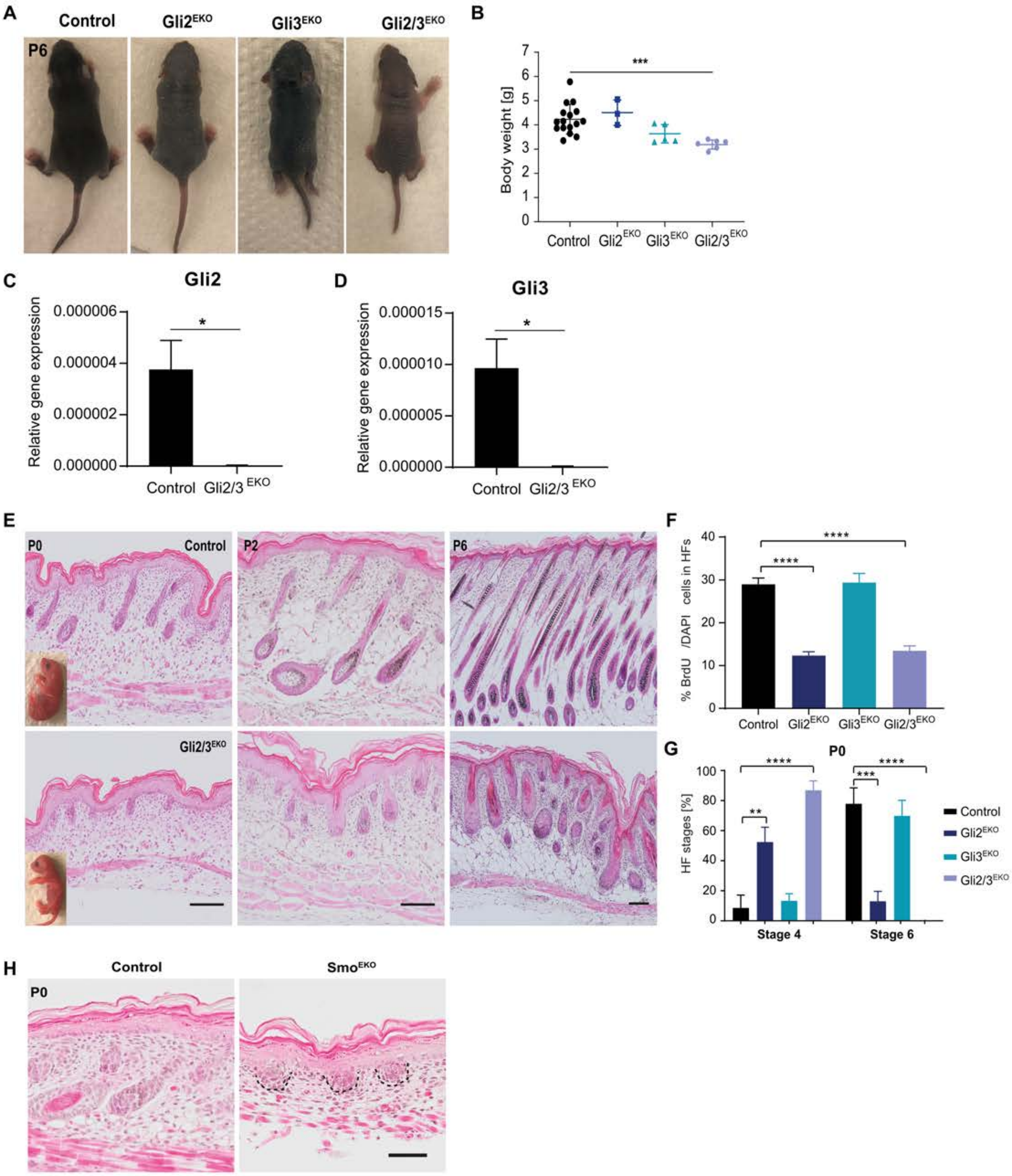


Supplementary Figure 1
Gozum et al.

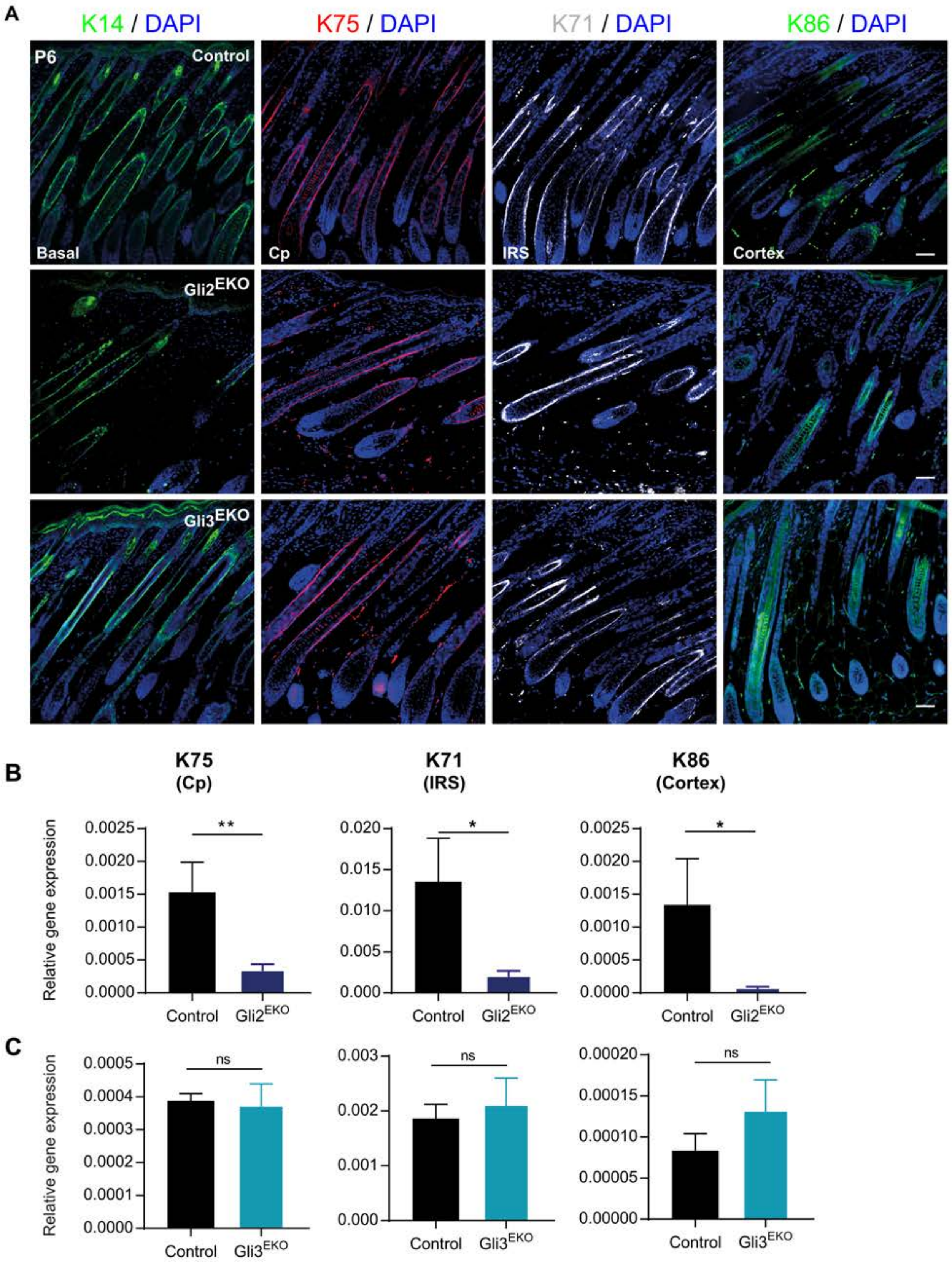


Supplementary Figure 2
Gozum et al.

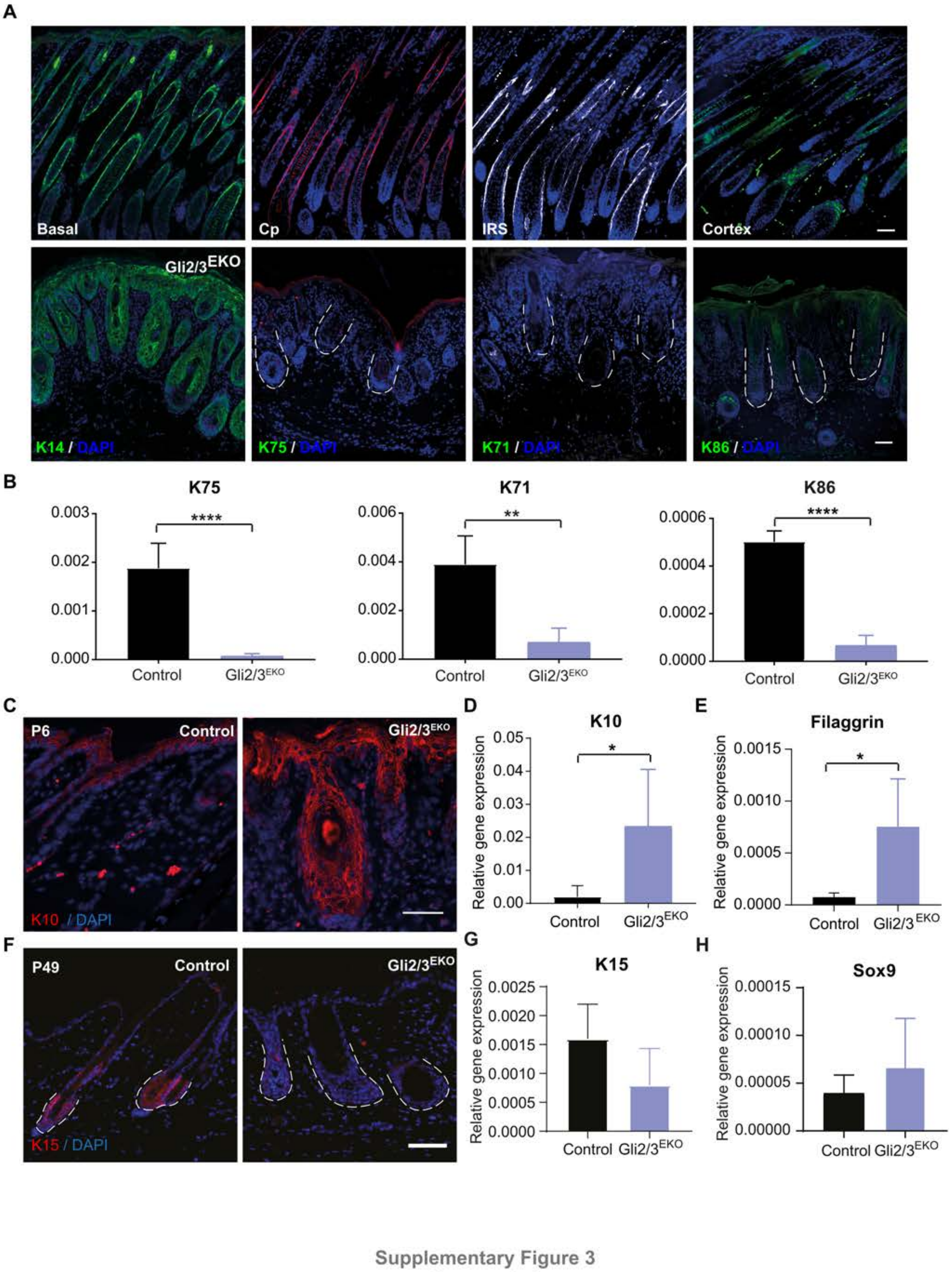


Supplementary Figure 3
Gozum et al

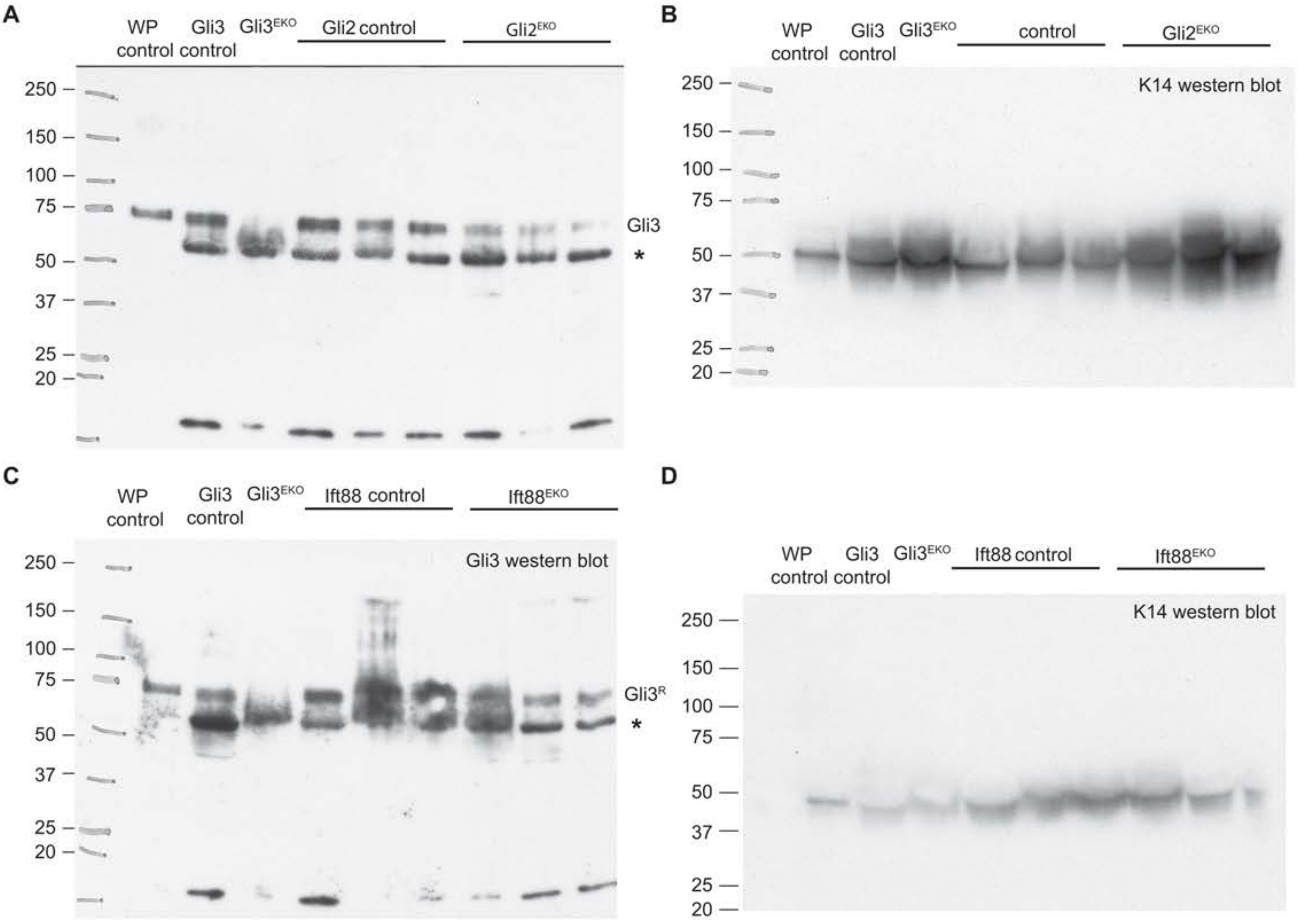


**Supplementary Figure 4**
**Gozum et al.**
